# The wheat *Sr22*, *Sr33*, *Sr35* and *Sr45* genes confer resistance against stem rust in barley

**DOI:** 10.1101/374637

**Authors:** M. Asyraf Md. Hatta, Ryan Johnson, Oadi Matny, Mark A. Smedley, Guotai Yu, Soma Chakraborty, Dhara Bhatt, Xiaodi Xia, Sanu Arora, Burkhard Steuernagel, Terese Richardson, Rohit Mago, Evans S. Lagudah, Nicola Patron, Mick Ayliffe, Matthew N. Rouse, Wendy A. Harwood, Sambasivam K. Periyannan, Brian J. Steffenson, Brande B. H. Wulff

## Abstract

In the last 20 years, stem rust caused by the fungus *Puccinia graminis* f. sp. *tritici* (*Pgt*), has re-emerged as a major threat to wheat and barley cultivation in Africa and Europe. In contrast to wheat with 82 designated stem rust (*Sr*) resistance genes, barley’s genetic variation for stem rust resistance is very narrow with only seven resistance genes genetically identified. Of these, only one locus consisting of two genes is effective against Ug99, a strain of *Pgt* which emerged in Uganda in 1999 and has since spread to much of East Africa and parts of the Middle East. The objective of this study was to assess the functionality, in barley, of cloned wheat *Sr* genes effective against Ug99. *Sr22*, *Sr33*, *Sr35* and *Sr45* were transformed into barley cv. Golden Promise using *Agrobacterium*-mediated transformation. All four genes were found to confer effective stem rust resistance. The barley transgenics remained susceptible to the barley leaf rust pathogen *Puccinia hordei*, indicating that the resistance conferred by these wheat *Sr* genes was specific for *Pgt*. Cloned *Sr* genes from wheat are therefore a potential source of resistance against wheat stem rust in barley.

## Introduction

Stem rust, caused by the fungus *P. graminis* f. sp. *tritici* (*Pgt*), is one of the major threats to barley (*Hordeum vulgare*) production in North America (Steffenson, 1992) and Australia (Dill-Macky et al., 1991). This destructive fungal disease can cause a significant reduction in plant growth and yield of both barley and wheat (De Wolf et al., 2011). In 1999, a new virulent isolate of *Pgt* called Ug99 (typed as race TTKSK according to Jin et al., 2008) was detected in Uganda which had overcome *Sr31*, a widely deployed stem rust resistance gene in bread wheat (*Triticum aestivum*) (Pretorius et al., 2000). At that time Ug99 and its derivatives were virulent on more than 80% of the world’s wheat cultivars (Singh et al., 2008). In recent years new *Pgt* races, that are not members of the Ug99 race group, have caused disease outbreaks on wheat in Europe (including Germany (Olivera Firpo et al., 2017), and Italy (Bhattacharya, 2017)), Asia (Russia (Shamanin et al., 2016)), and Africa (Ethiopia (Olivera et al., 2015)).

Effective ways of controlling this disease include fungicide application and breeding for resistant cultivars (McIntosh et al., 1995), with this latter strategy being the most cost effective and environmentally acceptable. However, when lines carrying a single resistance *(R)* gene effective against a specific disease are deployed, strong selection pressure is imposed on the pathogen population usually leading to resistance breaking down and the potential outbreak of an epidemic (Stakman, 1957). Notwithstanding, there are a few cases where *R* genes effective against *Pgt* have shown remarkable durability despite being deployed as a single gene for many years over a wide area where the pathogen is prevalent. Examples of such durability include *Sr31* which protected wheat from major losses for over 30 years until the Ug99 outbreak in 1999 (Ayliffe et al., 2008; Pretorius et al., 2000; Singh et al., 2006) and barley *Rpg1*, which has been widely deployed since the 1940s (Brueggeman et al., 2002). An alternative strategy is the simultaneous deployment of several *R* genes within a cultivar to prolong *R* gene efficacy in the field. There is no selective advantage for pathogen strains that have mutated to overcome a single *R* gene in the cultivar, thus imposing a barrier to the stepwise evolution of virulence (Dangl et al., 2013; Ellis et al., 2014; McDonald and Linde, 2002). However, it is difficult to ensure that multiple *R* genes, which may be scattered throughout the genome, remain together in a breeding program.

None-the-less, genetic resistance to cereal rust diseases has been fundamental for crop protection. For more than 100 years, breeders have introgressed resistance into wheat by undertaking wide crosses between wheat and its wild or domesticated relatives. Notable examples include the transfer of the stem rust resistance genes *Sr2* from emmer wheat (*Triticum turgidum* subsp. *dicoccum*) (McFadden, 1930), *Sr31*, *Sr50* and *Sr1RS^Amigo^* from rye (Mago et al., 2005b), *Sr24* and *Sr26* from *Thinopyrum ponticum* (Mago et al., 2005a), and *Sr36* from *T. timopheevi* (McIntosh and Gyarfas, 1971). However, sexual incompatibility and long generation times can impose significant barriers to successful gene introgression (Erickson, 1945). Also, linkage drag of deleterious alleles has hindered the deployment of many *Sr* genes in wheat, *i.e. Sr22* and *Sr43* due to yellow flour pigmentation and/or reduced yield and delayed heading date (Knott, 1984; Marais, 1992; Niu et al., 2014).

In contrast to wheat, where 82 stem rust resistance genes have been described (McIntosh et al., 2017), only seven stem rust resistance genes have been reported in barley; these being *Rpg1* (Brueggeman et al., 2002; Powers and Hines, 1933; Steffenson, 1992), *Rpg2* (Case et al., 2018; Patterson et al., 1957), *Rpg3* (Case et al., 2018; Jedel, 1990; Jedel et al., 1989), *rpg4* (Jin et al., 1994), *Rpg5* (Brueggeman et al., 2008; Sun and Steffenson, 2005; Sun et al., 1996), *rpg6* (Fetch et al., 2009) and *rpgBH* (Steffenson et al., 1984; Sun and Steffenson, 2005). *Rpg1* is the most widely deployed amongst these genes due to its broad-spectrum resistance which has remained effective for over 70 years (Brueggeman et al., 2002; Steffenson, 1992). However, a recent study showed that this gene is not effective to the Ug99 race TTKSK (Steffenson et al., 2017), leaving *rpg4/Rpg5* as the only gene complex known to confer resistance to this *Pgt* race in barley (Steffenson et al., 2009). In addition, seedling assays undertaken on two panels of 1,924 and 934 genetically diverse barley cultivars and wild barley accessions (*Hordeum vulgare* subsp. *spontaneum*), showed that more than 95% and 97% of accessions respectively, were susceptible to race TTKSK. Hence, it is important to identify novel sources of resistance to safeguard barley from stem rust (Steffenson et al., 2017). Given the limited number of *R* genes available for *Pgt* protection in barley, interspecies *R* gene transfer is a potentially valuable alternative (Wulff and Moscou, 2014).

The majority of *R* genes cloned encode proteins containing nucleotide-binding and leucine-rich repeat domains (NLR proteins) (Kourelis and van der Hoorn, 2018). Plant genomes typically contain several hundred NLR genes (Baggs et al., 2017). NLRs detect the presence of a pathogen by recognising pathogen effector molecules. This recognition can be direct, although more often it is indirect whereby the NLR (also known as the ‘guard’) recognises the effector-mediated modification of a host pathogenicity target, (also known as the ‘guardee’) (Dodds and Rathjen, 2010; Kourelis and van der Hoorn, 2018). NLR proteins that function by either mechanism have been successfully transferred by transgenesis to distantly related, nonsexually compatible species and shown to function in some instances. For example, the *L6* protein of flax (*Linum usitatissimum*, a member of the Linacea) directly binds a corresponding *AvrL567* effector protein of the flax rust pathogen *Melampsora lini*. When the *L6* gene is co-expressed with *AvrL567* in *Nicotiana benthamiana* (a member of the Solanaceae) a hypersensitive resistance response is activated (Dodds et al., 2004). Similarly, a number of *R* genes that function by guardee recognition have been shown to function upon interspecies transfer, exemplified by the transfer of the *Arabidopsis thaliana* (a Brassicaceae) guard and guardee gene pairs *RPS2* or *RPM1* with *RIN4* (Day et al., 2005; Chung et al., 2011) and *RPS5* with *PBS1* (Ade et al., 2007) into *N. benthamiana.*

Transferring *R* genes between species by conventional crossing can be a tedious task due to the extensive backcrossing usually required. However, it is now relatively straightforward to introduce these *R* genes as transgenes by transformation thereby avoiding this breeding requirement. Further advantages of transgenesis include that transfer is not limited to sexually compatible species, there is no linkage drag, and it becomes possible to stack multiple *R* genes at the same locus to ensure co-inheritance. When transferred between different species and families these *R* genes can function normally (reviewed in Wulff et al., 2011) and agronomically important examples include the *Bs2* gene from pepper (*Capsicum annuum*) which was successfully transferred to tomato (*Solanum lycopersicum*), another Solanacous species, where it confers resistance to bacterial leaf spot (Tai et al., 1999) and *CcRpp1* from pigeonpea (*Cajanus cajan*) which confers resistance to Asian soybean rust when introduced into soybean (*Glycine max*) (Kawashima et al., 2016).

Barley (*H. vulgare*) and wheat (*T. aestivum*) diverged from a common Triticeae ancestor approximately 10 to 14 million years ago (Schlegel, 2013) **(Figure 1a)**. It is therefore likely that wheat NLR genes will function in barley, and that wheat *Sr* genes could be used to improve the resistance of barley to *Pgt*. Nine major, dominant *Sr* genes have been cloned so far from wheat or its wild progenitors; these being *Sr13* from durum wheat (*T. turgidum* ssp. *durum*) (Zhang et al., 2017), *Sr21*, *Sr22* and *Sr35* from *T. boeoticum* and *T. monococcum* (Saintenac et al., 2013; Steuernagel et al., 2016,Chen et al., 2018), *Sr33*, *Sr45*, *Sr46*, and *SrTA1662* from *Aegilops tauschii* (Periyannan et al., 2013; Steuernagel et al., 2016; Arora et al., 2018), and *Sr50* from rye (*Secale cereale*) (Mago et al., 2015) **(Figure 1b)**. All these genes encode coiled-coil (CC)-NLR proteins and confer resistance to the Ug99 race group.

**Figure 1:**
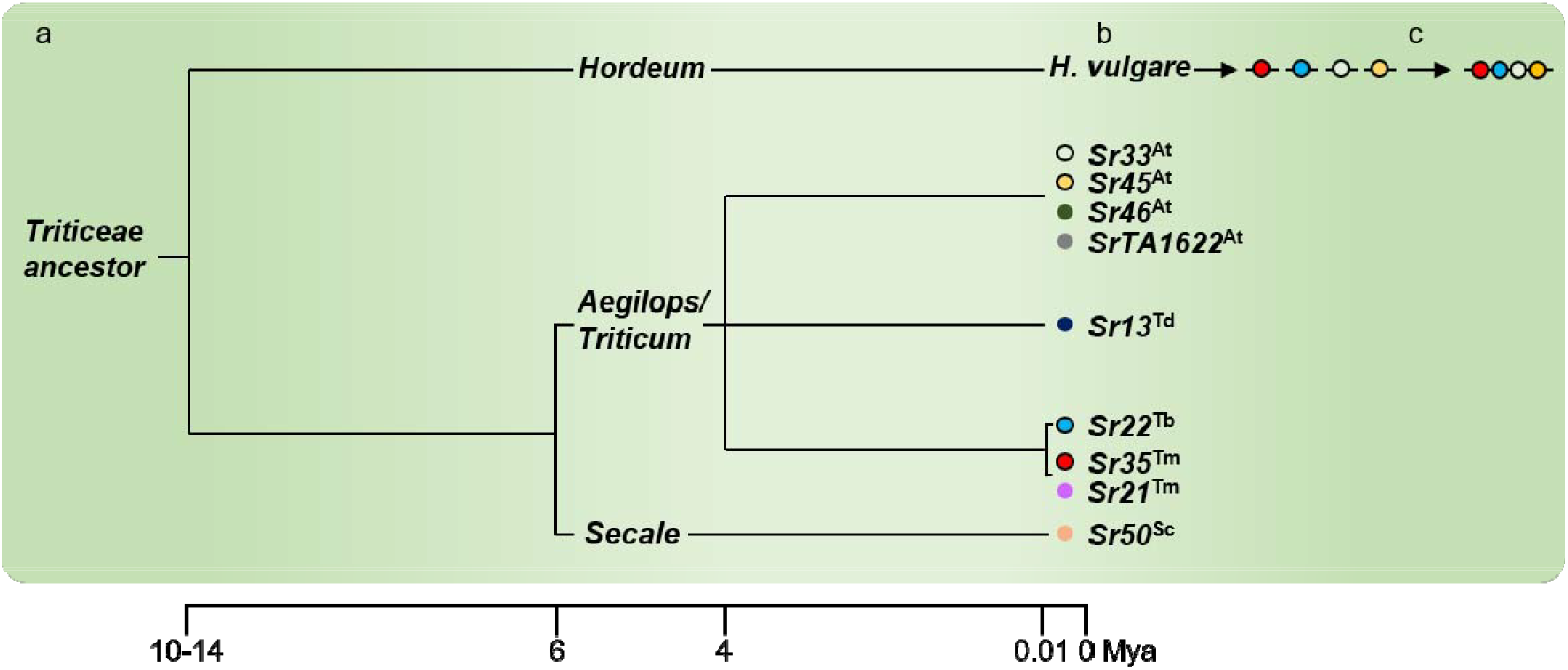
Strategy for improving barley resistance to stem rust with cloned wheat *Sr* genes. (a) Wheat and barley diverged from a common Triticeae ancestor 10-14 million years ago. (b) Cloned *Sr* genes from wheat, rye and the domesticated and wild relatives of wheat (coloured circles), function when transformed into barley (black outline, this study). (c) The future stacking of multiple cloned *Sr* genes in barley may provide durable resistance to wheat stem rust. At, *Aegilops tauschii*; Td, *Triticum turgidum* ssp. *durum*; Tb, *Triticum boeoticum*; Tm, *Triticum monococcum*; Sc, *Secale cereale*.

In the coming years, it is anticipated that there will be a large increase in the number of cloned *Sr* genes due to the development of rapid *R* gene isolation methods such as TACCA (Thind et al., 2017), MutRenSeq (Steuernagel et al., 2016), MutChromSeq (Sánchez-Martín et al., 2016), and AgRenSeq (Arora et al., 2018). Often the functional testing of *R* gene candidates is delayed by the need to isolate native regulatory sequences and to assemble large binary constructs encoding the *R* gene. This process can be accelerated by substituting regulatory elements from the previously cloned *R* genes, and generating *R* gene constructs using the type IIS restriction endonuclease-based Golden Gate cloning technique (Engler et al., 2008). Further, the incorporation of type IIS restriction sites allows the generation of user-defined overhangs thereby enabling simultaneous cloning of multiple fragments. This assembly method has dramatically decreased the amount of time required to design and develop gene constructs. However, one major requirement is that the fragments to be assembled must be free from recognition sites of the selected type IIS restriction endonuclease. This requires “sequence domestication” (removal of internal type IIS sites). While the open reading frame can be maintained due to the redundancy in the genetic code, the removal of sites from the regulatory sequences (introns, promoter and terminator) may affect gene expression and function.

In this study, we generated constructs encoding the wheat *Sr22*, *Sr33*, *Sr35* and *Sr45* genes using Golden Gate cloning and transformed these into barley **(Figure 1b)**. The resultant transgenic barley lines showed high-levels of resistance to *Pgt*. Future stacking of these *Sr* genes might therefore be used to engineer more durable immunity towards wheat stem rust in barley **(Figure 1c)**.

## Results

### The wheat *Sr22, Sr33, Sr35,* and *Sr45* genes confer resistance against wheat stem rust in transgenic barley

To determine whether cloned wheat *Sr* genes can function in barley to confer wheat stem rust resistance, barley cultivar (cv.) Golden Promise was transformed with constructs encoding either *Sr22* or *Sr33* via *Agrobacterium*-mediated transformation. The *Sr22* (9.8 kb) and *Sr33* (7.9 kb) sequences encoded their respective native 5’ and 3’ regulatory sequences and were identical in sequence to the endogenous wheat genes **(Figure 2a, 2b and Table S1**). Single-copy, hemizygous, primary transgenics were identified amongst T_0_ barley plants by Q-PCR so that single copy segregating T_1_ or T_2_ families, or homozygous T_2_ lines could be tested in subsequent generations.

**Figure 2:**
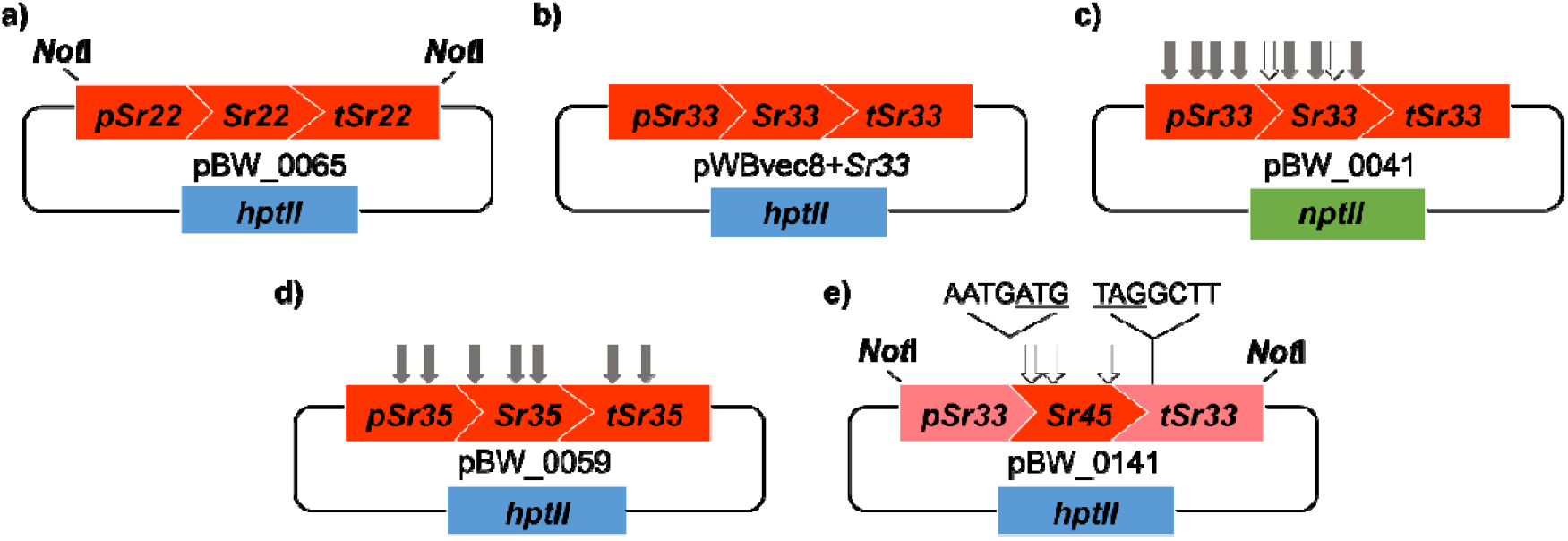
Schematic overview (not shown to scale) of the constructs described in this study. Binary construct containing (a,b) full-length *Sr22* and *Sr33*, respectively, driven by their 5’ and 3’ native regulatory elements, (c) *Bsa*I and *Bpi*I domesticated full-length *Sr33^d^* driven by its 5’ and 3’ native regulatory elements, (d) *Bpi*I domesticated full length *Sr35^d^* driven by its 5’ and 3’ native regulatory elements, and (e) *Bsa*I domesticated *Sr45^d^* driven by 5’ and 3’ *Sr33* regulatory elements. The non-native four-nucleotide linker in pBW_0141 was introduced immediately before the start codon (underlined) and immediately after the stop codon (underlined). Grey arrows correspond to the removed *Bpi*I sites and white arrows correspond to the removed *Bsa*I sites. Blue and green rectangles correspond respectively to *hpt*II and *npt*II plant selectable marker genes.

Eight progenies from ten *Sr22* segregating T_1_ families were inoculated with *Pgt* race MCCFC. Four families derived from lines 1370-11-01, 1370-17-01, 1370-19-01, 1372-08-01 segregated for resistance, while all the eight tested progenies from 1370-01-01 showed resistance **(Figure 3a and Table S2)**. The resistant individuals in these families showed near immunity to this *Pgt* isolate **(Figure 3a and Table S2)** whereas susceptible segregants and Golden Promise control seedlings all showed extensive *Pgt* growth. The presence of the *Sr22* transgene in resistant plants 1370-11-01 A, B, and C, and absence in susceptible sibling 1370-11-01-D and Golden Promise controls was confirmed by PCR amplification of the *hptII* selectable marker gene used for transformation **(Figure S1a)**. In contrast, the barley endogenous *CONSTANS* gene was amplified from all samples indicating that the lack of amplification of *hptII* from susceptible siblings and Golden Promise was not due to poor quality DNA **(Figure S2a)**.

**Figure 3:**
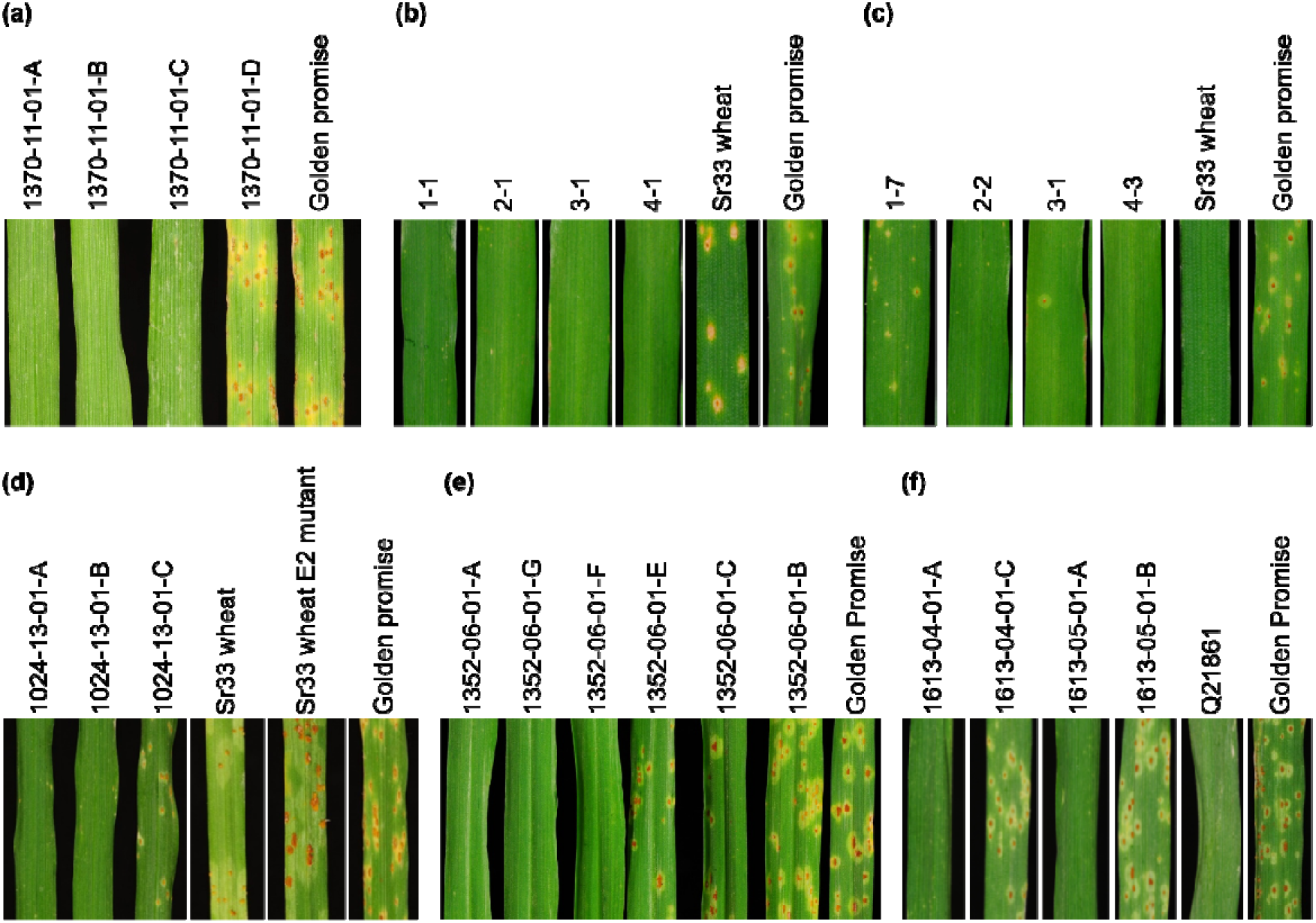
Transgenic *Sr22, Sr33, Sr33^d^, Sr35^d^,* and *Sr45^d^* expression provide resistance against stem rust in barley at the seedling stage. Stem rust infection assays using *Pgt* race (a) MCCFC on *Sr22* segregating T_1_ transgenics 1370-11-01 (A-D) and comparison to susceptible cv. Golden Promise wild type used in transformation, (b) MCCFC on *Sr33* T_2_ homozygous transgenics 1-1, 2-1, 3-1, 4-1 and comparison to resistant *Sr33* wheat and susceptible cv. Golden Promise, (c) TKTTF on *Sr33* T_2_ homozygous transgenics 1-7, 2-2, 3-1, 4-3 and comparison to resistant *Sr33* wheat and susceptible cv. Golden Promise, (d) MCCFC on *Sr33^d^* T_1_ segregating transgenics 1024-13-01 (A-C) and comparison to resistant *Sr33* wheat and the susceptible *Sr33* EMS-induced mutant wheat line (E2) and susceptible cv. Golden Promise, (e) TKTTF on T_2_ segregating *Sr35^d^* transgenics 1352-06-01 (A, B, C, E, F, G), and comparison to susceptible cv. Golden Promise, and (f) MCCFC on T_1_ segregating *Sr45* transgenics 1613-04-01 (A and C), 1613-05-01 (A and B) and comparison to resistant check Q21861 and susceptible cv. Golden Promise.

Four T_2_ lines, each derived from an independent transgenic event, were selected that were homozygous for the *Sr33* transgene and small T_3_ families (11-12 seedlings) from each line inoculated with either *Pgt* race MCCFC or TKTTF. All seedlings tested showed resistance to both races of *Pgt* **(Figure 3b and 3c and Table S3 and S4)**. In addition to susceptible Golden promise control seedlings, additional wheat control lines were included. The *Sr33*-containing cv. Chinese Spring and an EMS-derived mutant carrying a non-functional allele of *Sr33* (Periyannan et al., 2013) were used as resistant and susceptible controls, respectively, to demonstrate the avirulence of *Pgt* race MCCFC to *Sr33* **(Table S3)**.

### Wheat NLR genes can be sequence-modified but retain gene function

A functional wheat NLR transgene is typically 10 kb in length and consists of 4 kb of 5’ and 3’ regulatory elements, 3 kb of exons and 3 kb of introns. These long, contiguous sequences can be difficult to isolate and verify from a non-reference hexaploid wheat genome, and their synthesis is expensive. Multi-segment Golden Gate assembly (Weber et al., 2011) was therefore tested as an alternative for rapid and cost-effective generation of full-length *Sr* gene constructs using either native or non-native regulatory sequences. Firstly, the effect of sequence domestication (*i.e.* the removal of all Type IIS *Bsa*I and *Bpi*I restriction enzyme sites) was examined on *Sr33* function. Four *Bpi*I sites were removed from the *Sr33* promoter while 3 *Bpi*I and two *Bsa*I sites were removed from the *Sr33* open reading frame **(Figure 2c)**. Although the *Sr33* open reading frame was faithfully maintained through domestication, there was a risk that the removal of the four *Bpi*I sites in the 5’ regulatory sequence would disrupt gene function. The domesticated full-length *Sr33* gene (*Sr33^d^*) was transformed into cv. Golden Promise and single-copy primary transgenics again selected by Q-PCR. T_1_ progeny from 12 independent transgenic lines were infected with *Pgt* race MCCFC and nine T_1_ families shown to segregate for resistance while the remaining three families were all susceptible **(Figure 3d and Table S5).** PCR analysis of the *hptII* selectable marker gene again confirmed the presence of the transgene in resistant plants and its absence in susceptible plants **(Figure S1b)**, while the barley *CONSTANS* gene was amplified from all plant DNAs **(Figure S2b)**. These data confirm that *Sr33^d^* encodes functional *Sr33* resistance in spite of the sequence domestication process.

Having demonstrated that the endogenous wheat genes *Sr22* and *Sr33* can provide *Pgt* resistance in barley and that a sequence-modified version of *Sr33* (*Sr33^d^*) maintains function, two additional domesticated wheat *Sr* genes were developed. A domesticated *Sr35* construct (*Sr35^d^*) was generated by multi-segment Golden Gate assembly which involved removing seven *Bpi*I sequences from the gene **(Figure 2d)**. Three T_2_ families, each derived from an independent *Sr35^d^* barley transgenic, were tested with *Pgt* race TKTTF and each segregated for resistance **(Figure 3e and Table S6)**. A chimeric *Sr45* gene construct (*Sr45^d^*) was also assembled by Golden Gate and consisted of a *Bsa*I domesticated *Sr45* open reading frame flanked by *Sr33* 5’ and 3’ regulatory sequences **(Figure 2e)**. This construct did not require the removal of *Bpi*I sites from *Sr33* regulatory sequences, however the assembly resulted in the introduction of four additional nucleotides at both the junction between the *Sr33* promoter and start codon of *Sr45* and the termination codon of the *Sr45* ORF and the *Sr33* 3’ regulatory sequence **(Figure 2e)**. In spite of these modifications four out of eleven *Sr45^d^* T_1_ families, each derived from an independent primary transgenic line, segregated for *Pgt* resistance **(Figure 3f and Table S7).** For these latter two transgenes their presence in resistant plants and absence in susceptible siblings and control seedlings was again confirmed using *hptII* and *CONSTANS* PCR analyses (**Figure S1c, S1d, S2c and S2d)**. These data demonstrate that these wheat NLR genes can be sequence-modified to facilitate further molecular biological manipulation and that regulatory sequences can be functionally exchanged between NLR genes in some instances.

### Pathogen and race-specific resistance is maintained by wheat *Sr* genes in barley

To rule out the possibility that these resistant transgenic barley lines are a consequence of an ectopic non-specific defence reaction, we tested these transgenic plants with the barley leaf rust pathogen. All *Sr22, Sr33, Sr33^d^, Sr35^d^* and *Sr45^d^* transgenic barley lines, as well as Golden Promise were susceptible to *P. hordei*. In contrast, a barley control, accession PI531901-4, and the wheat *Sr33* line, which is a nonhost of *P. hordei*, were both resistant indicating that the barley resistance observed in stem rust infection assays was specific to *Pgt* **(Figure S3 and Table S8, S9, S10, S11, and S12)**. Ten *Sr35^d^* transgenic T_1_ families were also tested with *Pgt* race MCCFC (virulent to *Sr35*) including the three T_2_ lines described above that are resistant to *Pgt* race TKTTF. All ten *Sr35^d^* T_1_ families were susceptible to *Pgt* race MCCFC (**Table S13**) indicating that race-specificity of this wheat gene is maintained in transgenic barley.

## Discussion

Barley is a major food staple in the mountainous areas of Central Asia, Southwest Asia, and Northern Africa (Von Bothmer et al., 2003). The re-emergence of wheat stem rust as a major biotic constraint to wheat production also poses a threat to barley production. A recent study revealed very limited resistance to *Pgt* isolate Ug99 in both cultivated barley and its immediate progenitor *H. vulgare* ssp. *spontaneum* (Steffenson et al., 2017). This *Pgt* isolate has caused major epidemics in East Africa since 1999. One avenue for improving resistance to stem rust in barley is to utilise diverse genetic resistance from outside the barley gene pool.

*R* genes typically function when transferred from one species to another within the same family (Wulff et al., 2011). In this study, the wheat *Sr22, Sr33, Sr35,* and *Sr45* genes were shown to function when transferred to barley and confer race-specific disease resistance to *Pgt*. Other examples of *R* gene transfer in monocots include the introduction of the maize NLR gene *Rxo1* into rice where it confers resistance to bacterial streak disease (Zhao et al., 2005) and single-cell transient expression assays of the barley *Mla6* gene in wheat where it confers *AvrMla6*-dependent resistance to *Blumeria graminis* f. sp. *hordei* (*Bgh)* (Halterman et al., 2001). In concordance, the wheat *Sr22*, *Sr33, Sr35,* and *Sr45* genes also function in barley, suggesting that the downstream signalling pathway(s) of NLR proteins in wheat and barley has remained conserved since the divergence of these two species 10 to 14 million years ago (Schlegel, 2013).

The functional transfer of these wheat *Sr* genes into barley potentially provides additional sources of stem rust resistance in this recipient species. Interestingly, in most cases, these barley transformants displayed a highly resistant reaction that is stronger than that observed for the endogenous wheat genes. Similarly, when the barley *Rpg1* gene was expressed as a transgene in barley, this also gave rise to a near-immune reaction (Horvath et al., 2003). In contrast to these transgenic barley experiments, near-immune reactions were extremely rare when large scale screening of wild and cultivated barley lines was undertaken using different *Pgt* races (e.g. Steffenson et al., 2017). The increased resistance conferred by these transgenes may be a consequence of elevated expression arising from position effects or alternatively their interaction with a new genetic background in the case of interspecies transfer.

Importantly we confirmed that race specificity of the *Sr35* gene was maintained in barley. We assume this is also the case for *Sr22*, *Sr33* and *Sr45*, but cannot confirm this due to an absence of *Pgt* races virulent on Golden Promise and avirulent to these genes. Unlike most barley cultivars, Golden Promise does show a high level of resistance to many *Pgt* races which makes *Sr* gene analysis difficult as it is one of the few transformable barley cultivars available. However, for these latter genes we confirmed that resistance is not a consequence of a non-specific defence reaction caused by ectopic expression of these genes. All transgenic barley lines tested with *P. hordei*, the causal agent of barley leaf rust, were as susceptible as Golden promise control lines demonstrating species-specific resistance conferred by these *Sr* genes. The phylogenetic relatedness and very similar lifecycle of these two fungal pathogen species argues against a generic defence response being activated. Interestingly these data also suggest that there is little conservation of effectors recognised by these wheat *Sr* genes in *P. hordei*.

In the last couple of years, many significant improvements have been made in the field of *R* gene cloning. For example, sequence comparison of multiple independently-derived mutants, facilitated by various genome complexity reduction technologies, e.g. NLR exome capture (Steuernagel et al., 2016) or chromosome flow sorting (Sánchez-Martín et al., 2016,Thind et al., 2017) was used to rapidly clone *Sr22*, *Sr45, Pm2* and *Lr22a* from hexaploid wheat. Recently, the requirement of mutagenesis was overcome by combining association genetics with NLR exome capture on a diversity panel of *Ae. tauschii* (Arora et al., 2018). The resulting application, AgRenSeq, allowed the rapid cloning of *Sr46* and *SrTA1662* (Arora et al., 2018). These advances coupled with the recent availability of a wheat reference genome will greatly accelerate *R* gene discovery and cloning.

As more wheat *R* genes are cloned, they can be tested in barley using the strategy demonstrated in this paper and, in the case of stem rust potentially provide greater control of stem rust disease in this species. The ability to modify these *R* gene sequences by multi-segment Golden Gate assembly (Weber et al., 2011), even including regulatory elements, and yet maintain gene function will greatly facilitate the manipulation and validation of these genes. Unlike hexaploid wheat, the diploid nature of barley will help understand the fundamental aspects of wheat stem rust resistance. Its greater amenability to mutagenesis will enable the identification of additional genes required for rust *R* gene function, as well as potential host susceptibility genes. Interestingly a higher proportion of rust *R* genes in barley are recessive (26.3%) compared to wheat (6.7%) (Uauy et al., 2017) and the cloning of recessive resistance genes may provide novel fundamental insight into plant pathogen interactions.

NLR genes are not the sole means of generating disease resistant plants. Another approach to improve wheat stem rust resistance in barley is to combine multiple, additive minor effect quantitative trait loci (QTLs). Bi-parental and genome-wide association studies (GWAS) have identified QTLs associated with stem rust resistance in barley (Mamo, 2013; Sallam et al., 2017; Turuspekov et al., 2016; Zhou et al., 2014; Case et al., 2018). More recently, GWAS on adult plants identified seven novel QTLs conferring adult plant resistance to *Pgt* race QCCJB and a mixed inoculum of races TTKSK, TTKST, TTKTK and TTKTT, which are all members of the Ug99 group (Case et al., 2017). The presence of APR genes or minor effect QTLs has been shown to enhance the strength of race-specific *R* genes (Hiebert et al., 2016) and promote their longevity (Brun et al., 2010).

Interestingly an APR gene has also been transferred between monocot species by transgenesis and shown to function. The wheat *Lr34* adult plant resistance gene (APR), which encodes an ABC transporter, has been shown to provide resistance against multiple, diverse rust, mildew and blast fungal pathogens in barley, rice, durum wheat, maize and sorghum (Risk et al., 2013; Krattinger et al., 2016; Rinaldo et al., 2017; Sucher et al., 2017; Schnippenkoetter et al., 2017), although the mechanism of this resistance is as yet unknown. Wide interspecies transfer of functional disease resistance is therefore not limited to NLR genes.

Given that wheat *R* genes function in barley, it is reasonable to expect that barley *R* genes will also function in wheat. Therefore, barley *R* genes conferring resistance to wheat stripe rust (*P. striiformis* f. sp. *tritici*) (Dawson et al., 2016) might be deployed in wheat to control this disease. However, caution should be taken so that *R* genes transferred from one crop to another are not easily overcome which would potentially facilitate a host jump and create a new disease problem. Ideally, *R* genes with different specificities should be combined as a multi *R* gene stack, preferably with the inclusion of APR genes **(Figure 1c)**. This is likely to confer more durable resistance by delaying the emergence of resistance-breaking strains of the pathogen (Dangl et al., 2013; McDonald and Linde, 2002).

In summary, functional transfer of the *Sr22*, *Sr33, Sr35* and *Sr45* genes into barley has created a new source of resistance to stem rust in barley. As more novel rust *R* genes are cloned and shown to function in barley, these could subsequently be deployed in a stack to provide broad-spectrum resistance and reduce the risk of resistance breakdown. Future GM field experiments with barley plants expressing single or multiple *Sr* transgenes will be useful to assess the agronomic value of wheat *Sr* genes in barley cultivation.

## Experimental procedures

### Generation of binary constructs carrying *Sr* genes

To assemble a plant transformation construct containing an *Sr22* expression cassette, a 9,855 bp fragment of DNA containing the *Sr22* coding sequence, 2,377 bp of 5’ regulatory sequence (*i.e.* 5’ of the predicted start codon) and 1,560 bp of 3’ regulatory sequence (*i.e*. 3’ of the STOP codon) was synthesised by a commercial DNA synthesis provider (Life Technologies Ltd) with flanking *Not*I sites. The synthetic DNA was cloned into the *Not*I site of the pVec8 binary vector (Wang et al., 1998).

The *Sr33* gene sequence from the binary vector pVecNeo+*Sr33* (Periyannan et al. 2013) was introduced into the binary vector pWBvec8 (Steuernagel et al. 2016) using *Psp*OMI and *Not*I restriction enzymes. The *Sr33* gene sequence in the later construct pWBvec8+*Sr33* was proof read using AtM5F1, AtM5R1, AtM5F2, AtM5R2, AtM5F3 and AtM5R3 primers (Periyannan et al. 2013).

To generate *Sr33^d^*, a 7,854 bp fragment of *Sr33,* including 2,381 bp of 5’ and 1,405 bp of 3’ regulatory sequence and an 8,255 bp fragment of *Sr35^d^*, including 2,462 bp of 5’ and 2,615 bp of 3’ regulatory sequence was synthesised flanked by a pair of divergent *Bpi*I recognition sites. Prior to synthesis, any recognition sequences for the restriction endonucleases *Bsa*I and *Bpi*I were removed by introducing synonymous mutations in coding sequences and avoiding intron splice junctions. This fragment was simultaneously assembled using Golden Gate cloning (Weber et al., 2011) into a level two Golden Gate acceptor plasmid, pAGM4723 (Weber et al., 2011), with a hygromycin selectable marker cassette to confer resistance to hygromycin.

A transformation construct for barley containing an *Sr45^d^* expression cassette was constructed as described in Arora et al., (2018)and the assembled gene cassette was cloned into the *Not*I site of the pVec8 binary vector. All binary plasmids containing the desired insert were transformed into *Agrobacterium tumefaciens* (strain AGL1) for transformation of barley.

### Barley transformation

*Agrobacterium*-mediated transformation of *Sr* gene constructs *Sr22*, *Sr33^d^*, *Sr35^d^* and *Sr45^d^* into barley cv. Golden Promise was performed as described in Harwood (2014). Ten to 12 independent primary transgenic (T_1_) plants carrying the *Sr* gene construct were recovered. Confirmation that the transformants carried the *Sr* gene was done by PCR on genomic DNA using gene specific markers **(Table S14)**. The copy number analysis of *Sr22*, *Sr33^d^*, *Sr35^d^*, and *Sr45^d^* by Q-PCR was outsourced to iDNA Genetics, Norwich Research Park, UK. Plants with a single copy transgene were selected and propagated for phenotyping. Binary vector pWBvec8+*Sr33* was transformed into cv. Golden Promise using *Agrobacterium*-mediated transformation as described in Moore et al. (2015). Four advanced generation lines, SH1, SH2, SH3 and SH4, were selected as homozygous for the *Sr33* transgene by screening with *Sr33* sequence-specific primers.

### Functional testing of *Sr33* and *Sr45* with *Pgt* race MCCFC

To identify a *Pgt* race which would be virulent on Golden Promise and avirulent on *Sr33* and *Sr45*, we interrogated a panel of 151 *Ae. tauschii* accessions which had their NLR repertoires sequenced (Arora et al., 2018) by BLAST search with the *Sr33*, *Sr45*, *Sr46* and *SrTA1662* gene sequences (using a ≥99% identity and 100% query coverage cut-off). We identified one accession that appeared to contain only *Sr33*, and four accessions that appeared to contain only *Sr45*. These accessions were resistant to *Pgt* race MCCFC, a race which was previously shown to be virulent on Golden Promise (Arora et al., 2013; Kleinhofs et al., 2009), whereas 31 *Ae. tauschii* accessions which did not appear to contain any of the aforementioned *Sr* genes were predominantly susceptible or intermediate in their response to MCCFC (Table S15). From this we concluded that MCCFC is avirulent towards *Sr33* and *Sr45*.

### Wheat rust inoculations and phenotypic evaluations

For the stem rust inoculations, *Sr* barley (*Sr22, Sr33^d^, Sr35^d^, Sr45^d^*) T_1_ or T_2_ plants alongside with the susceptible control cv. Golden Promise were infected with *Pgt* race MCCFC or/and TKTTF 10 days after planting. The inoculated plants were rated for disease response 12-14 days after inoculation as previously described in Yu et al., (2017). For *Sr33,* T_2_ plants alongside with the susceptible control cv. Golden Promise were infected with *Pgt* race MCCFC (isolate 59KS19) and TKTTF (isolate 13ETH18-1) and the inoculation, incubation and disease assessment procedures were performed as described previously (Zhang et al., 2017). At least 10 plants from each homozygous family were evaluted for *Sr33* and infection types were recorded once, 12 days after inoculation.

For the leaf rust experiment, each cone rack (98 cones, three seeds per cone, so in total 294 plants per cone rack) received 1 ml of inoculum (15 mg spore) of *P. hordei* race 4 (Levine and Cherewick, 1952) across the primary leaves of 8-9 day-old seedlings. *P. hordei* isolate 12TX15-2 was used to inoculate the *Sr33-*transformed lines. Therefore, each plant received 0.05 mg of urediniospores. To minimize risk of phytotoxicity, the Soltrol 170 oil carrier was evaporated from the leaf surfaces by two hours of gentle fanning under 400-watt HPS light bulbs. Inoculated seedlings were incubated at 22 °C inside mist chambers with a 100% relative air humidity provided by a household ultrasonic humidifier. Post inoculation, plants were moved to a greenhouse running a 16-hour day length with a night temperature of 15 °C and a day temperature of 20 °C. The information of the resistant control used in the barley leaf rust infection assays, PI531901-4 can be accessed via this link: https://npgsweb.ars-grin.gov/gringlobal/accessiondetail.aspx?id=1426837.

Disease phenotypes (*i.e.* infection types (ITs) were scored twice for each experiment: first at 10 and then at 12 days post inoculation (dpi) as previously described in Park et al., (2017).

### PCR primers and amplification

PCR assay was performed to confirm the presence of the transgene in transgenic plants or the ability to PCR-amplify the control, the endogenous *CONSTANS* gene. Specific PCR primers for each (trans)gene were designed using the web-based application Primer3 (http://primer3.ut.ee/) **(Table S14)**. PCRs with a final volume of 20 μl contained 10 ng of genomic DNA, 10 μl of REDTaq ReadyMix PCR Reaction Mix (Sigma-Aldrich, St Louis, MO, USA) and 10 μM of each primer. The reaction schedule for each transgene was; Neomycin phosphotransferase II (*nptII*) gene 94 °C for 5 min, 35 cycles of 94 °C for 30 s, 60 °C for 40 s and 72 °C for 70 s, 72 °C for 10 min and 16 °C; Hygromycin phosphotransferase II (*hptII*) gene 95 °C for 5 min, 29 cycles of 94°C for 30 s, 54 °C for 30 s and 72°C for 30 s, and 16 °C; *Sr22* and *Sr45* genes 95 °C for 5 min, 29 cycles of 94 °C for 30 s, 59 °C for 30 s and 72 °C for 30 s, and 16 °C. The reaction schedule for the *CONSTANS* gene was; 95 °C for 5 min, 33 cycles of 94 °C for 30 s, 57 °C for 30 s and 72 °C for 30 s, and 16 °C.

## Acknowledgements

The authors wish to thank the support of the Two Blades Foundation, USA, the Lieberman-Okinow Endowment at the University of Minnesota, the USDA-ARS, and the Biotechnology and Biological Sciences Research Council, UK, Grant numbers BB/H019820/1, BB/L009293/1 and BB/L011794/1. MAMdH was supported by a fellowship from Universiti Putra Malaysia (UPM), Malaysia. The authors declare no conflict of interest.

## Author contributions

MAMdH, MS, RM, GY, NP, and MA designed and generated *Sr22*, *Sr33^d^*, *Sr35^d^* and *Sr45^d^* constructs. MAMdH, MA and WH performed *Sr22*, *Sr33^d^*, *Sr35^d^* and *Sr45^d^* transformation. RJ, OM, MR and BJS phenotyped *Sr22*, *Sr33^d^*, *Sr35^d^* and *Sr45^d^* transgenics. SC, DB, XX, TR, RM and MA generated the *Sr33* construct, performed transformation and selected homozygous lines while MNR phenotyped the transgenics. SA and BS performed *Ae. tauschii* sequence analysis. BBHW, SKP, BJS, EL, NP and WH conceived and designed study. MAMdH drafted manuscript with input from BBHW, BJS, MA, SP and NP. All authors read and approved the final manuscript.

## Supporting information

**Figure S1** Resistance to wheat stem rust in *Sr22, Sr33^d^, Sr35^d^,* and *Sr45^d^* barley transgenic T_1_ or T_2_ families segregating for the transgene correlates with the presence of the transgene.

**Figure S2** Amplification of the barley endogenous CONSTANS gene using genomic DNA of *Sr* gene transgenics and Golden Promise (positive control) as a template.

**Figure S3** Leaf rust infection assays with *P. hordei* race 4 on *Sr22*, *Sr33*, *Sr33^d^*, *Sr35^d^*, and *Sr45^d^* representative T_1_ and T_2_ transgenics at the seedling stage.

**Table S1** List of binary constructs carrying *Sr* gene.

**Table S2** Stem rust infection assays with *Pgt* race MCCFC on *Sr22* T_1_ families.

**Table S3** Stem rust infection assays with *Pgt* race MCCFC on *Sr33* T_2_ homozygous lines.

**Table S4** Stem rust infection assays with *Pgt* race TKTTF on *Sr33* T_2_ homozygous lines.

**Table S5** Stem rust infection assays with *Pgt* race MCCFC on *Sr33^d^* T_1_ families.

**Table S6** Stem rust infection assays with *Pgt* race TKTTF on *Sr35^d^* T_2_ families. **Table S7** Stem rust infection assays with *Pgt* race MCCFC on *Sr45^d^* T_1_ families. **Table S8** *Puccinia hordei* race 4 infection assays on *Sr22* T_2_ families.

**Table S9** *Puccinia hordei* race 4 infection assays on *Sr33* T_2_ homozygous lines. **Table S10** *Puccinia hordei* race 4 infection assays with on *Sr33^d^* T_2_ families. **Table S11** *Puccinia hordei* race 4 infection assays on *Sr35^d^* T_2_ families.

**Table S12** *Puccinia hordei* race 4 infection assays on *Sr45^d^* T_2_ families.

**Table S13** Stem rust infection assays with *Pgt* race MCCFC on *Sr35* T_1_ families.

**Table S14** PCR primers used in this study.

**Table S15** Functional testing of *Sr33* and *Sr45* with *Pgt* race MCCFC.

